# Cell-free expressed membraneless organelles sequester RNA in synthetic cells

**DOI:** 10.1101/2023.04.03.535479

**Authors:** Abbey O Robinson, Jessica Lee, Anders Cameron, Christine D Keating, Katarzyna P. Adamala

**Affiliations:** Department of Genetics, Cell Biology and Development, University of Minnesota, Minneapolis, MN 55455, USA; Department of Chemistry, The Pennsylvania State University, University Park, PA 16802, USA

## Abstract

Compartments within living cells create specialized microenvironments, allowing for multiple reactions to be carried out simultaneously and efficiently. While some organelles are bound by a lipid bilayer, others are formed by liquid-liquid phase separation, such as P-granules and nucleoli. Synthetic minimal cells have been widely used to study many natural processes, including organelle formation. Here we describe a synthetic cell expressing RGG-GFP-RGG, a phase-separating protein derived from LAF-1 RGG domains, to form artificial membraneless organelles that can sequester RNA and reduce protein expression. We create complex microenvironments within synthetic cell cytoplasm and introduce a tool to modulate protein expression in synthetic cells. Engineering of compartments within synthetic cells furthers understanding of evolution and function of natural organelles, as well as it facilitates the creation of more complex and multifaceted synthetic life-like systems.

## Main

The appearance of discrete intracellular compartments was critical for the evolution of eukaryotes. On average, eukaryotic cells are larger and functionally more complex than prokaryotic cells. This complexity can be attributed to the existence of the nucleus, as well as other compartments and organelles. The self-assembly of and spatiotemporal organization provided by intracellular compartments facilitates simultaneous and concatenate cell-processes that are unattainable in simpler organisms^1^.

A key goal in bottom-up synthetic biology is to build artificial cells that mimic the functions and organization of living cells to create complex programmable bioreactors for foundational research and many practical applications^2^. To achieve this goal, there have been numerous efforts to build organelle-like compartments within synthetic cells to increase the complexity of synthetic cells and to better model and understand intracellular environments in natural cells. These efforts include engineering proto-organelles based on natural endosymbionts^3–6^, oligolamellar vesicles^7,8^, and lipid sponge droplets^9^. Inspired by membraneless organelles present in living cells, there has also been work on engineering protein-rich liquid droplets called coacervates as membraneless organelles for artificial cells^10–13^.

Membraneless organelles are cellular compartments that lack a lipid bilayer. They are formed by liquid-liquid phase separation (LLPS), a thermodynamic-driven event characterized by the de-mixing of liquid phases—in cells, usually a dense protein- and/or RNA-rich phase separates from more dilute components. Many of these membraneless condensates are formed by intrinsically disordered proteins (IDPs): dynamic proteins that lack tertiary structure. IDPs are ubiquitous, with approximately 30% of human proteins containing intrinsically disordered regions^14,15^. Ribonucleoprotein (RNP) particles are a type of these intracellular condensates formed between RNA and RNA-binding proteins which create cellular structures such as P-granules, stress granules, and nucleoli^16–19^. While RNP particles have diverse functions, many are involved in RNA processing, degradation, and storage, thus exerting a level of control over gene expression^20,21^.

LAF-1 is an RNA helicase found in *Caenorhabditis elegans* P-granules which contain an intrinsically disordered region called the RGG (arginine-glycine-glycine) domain. Previously, it was found that RGG domains are critical for phase separation of LAF-1, and that the RGG domains themselves undergo liquid-liquid phase separation^22^. The RGG domain has low sequence complexity and consists of multiple arginine-glycine-glycine repeats. RGG domains often occur in repeats and are conserved in many other phase separating proteins^23^. RGG domains are also known to bind RNA, including G-quadraplexes^21,24–26^.

More recently, the RGG domain was isolated and engineered to function as a synthetic membraneless organelle for both living and protocells^27^. Engineering synthetic cells that mimic the complexity and organization of living cells is a key goal in synthetic biology. The creation of compartments within liposome-based synthetic cells, or, better yet, a synthetic cell that can create its own compartments through cell-free gene expression will allow for more orthogonal capabilities of these artificial cells.

Significant obstacles to creating membrane-bound organelles within synthetic cells include issues with fusion of outer and inner vesicles, regulating the permeability of organelle membranes, and lipogenesis^28^. Thus, creation of artificial organelles that are membraneless is an attractive option. More specifically, a major goal in pursuit of creating robust synthetic cells is regulation of gene expression, and membraneless organelles are known to sequester RNA, thereby playing a role in protein expression^21^.

Many biological tools for modulation of gene expression that exist in complex living organisms have not been reconstituted in synthetic cells^29^. Here, we report the creation of an artificial membraneless organelle that can be expressed cell-free inside of synthetic cells and can sequester RNA to modulate protein expression (Fig. 1). We find that RGG-GFP-RGG, a recombinant protein with two RGG domains flanking GFP, forms coacervates when expressed by *in vitro* transcription and translation (TXTL), both in solution and when encapsulated inside of liposomes^30–32^. We observe localization of fluorescently labeled polyU RNA to cell-free expressed RGG-GFP-RGG. We also find that the presence of RGG-GFP-RGG in TXTL reactions expressing another plasmid-encoded fluorescent protein sequesters RNA produced during transcription and results in reduced expression of protein.

**Fig. 1:**
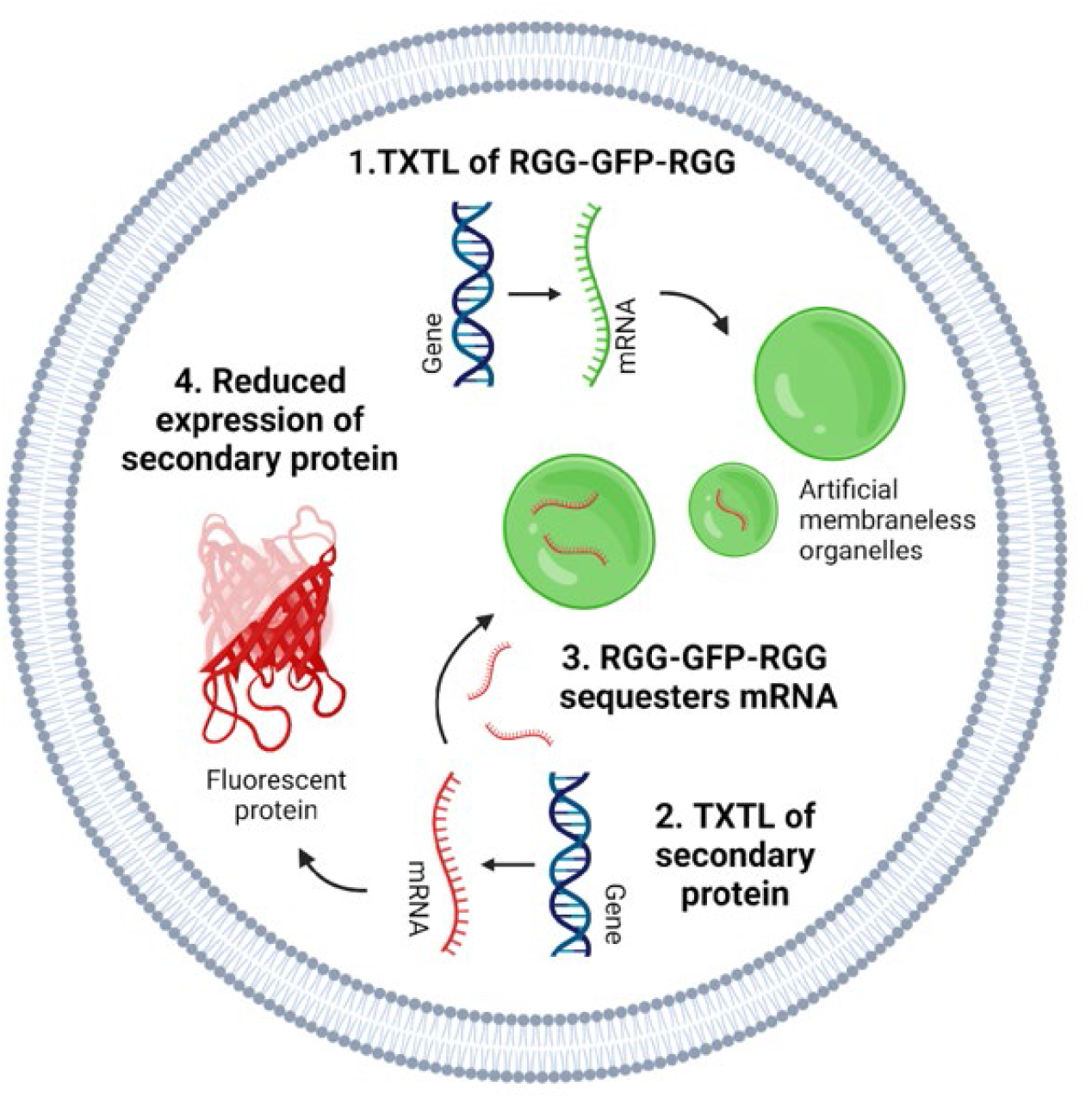
Schematic illustration of membraneless organelle expression and function in synthetic cell. A plasmid encoding for RGG-GFP-RGG, a phase separating protein, is expressed by TXTL to create coacervate droplets (membraneless organelle) inside of a liposome. When another plasmid—encoding for second fluorescent protein—is expressed by TXTL, RGG-GFP-RGG sequesters transcriptional mRNA, reducing expression and fluorescence of the secondary protein.

## Results

### Cell-free expression of RGG-GFP-RGG forms coacervates inside of liposomes

We first determined if a coacervate-forming protein could be expressed by *in vitro* transcription and translation (TXTL). RGG-GFP-RGG, an intrinsically disordered protein labeled with green fluorescent protein, was used for cell-free protein expression experiments. The RGG protein was chosen for its relatively heterogeneous amino acid sequence, compared to other coacervate-forming proteins, as we speculated highly repetitive sequences may be less compatible with TXTL expression. RGG-GFP-RGG contains two RGG domains flanking GFP, allowing for quantitative fluorescence spectroscopy and microscopy. RGG-GFP-RGG demonstrates upper critical solution temperature (UCST) phase behavior, meaning that above the critical temperature the proteins are miscible in solution, and below the critical temperature form coacervates^27^. In physiological buffer (150mM NaCl, pH 7.5) the critical temperature of RGG-GFP-RGG is around 40°C^27^. RGG-GFP-RGG forms liquid-like, spherical droplets at ambient temperatures and is reversibly miscible upon heating and cooling^27^. Previously, phase separation by purified RGG-GFP-RGG was observed in protocell-like structures formed by aqueous droplets in oil^27^. For control experiments, RGG-GFP-RGG was purified from *E. coli* according to previous methods and phase separation was observed (Fig. 2a).

**Fig. 2:**
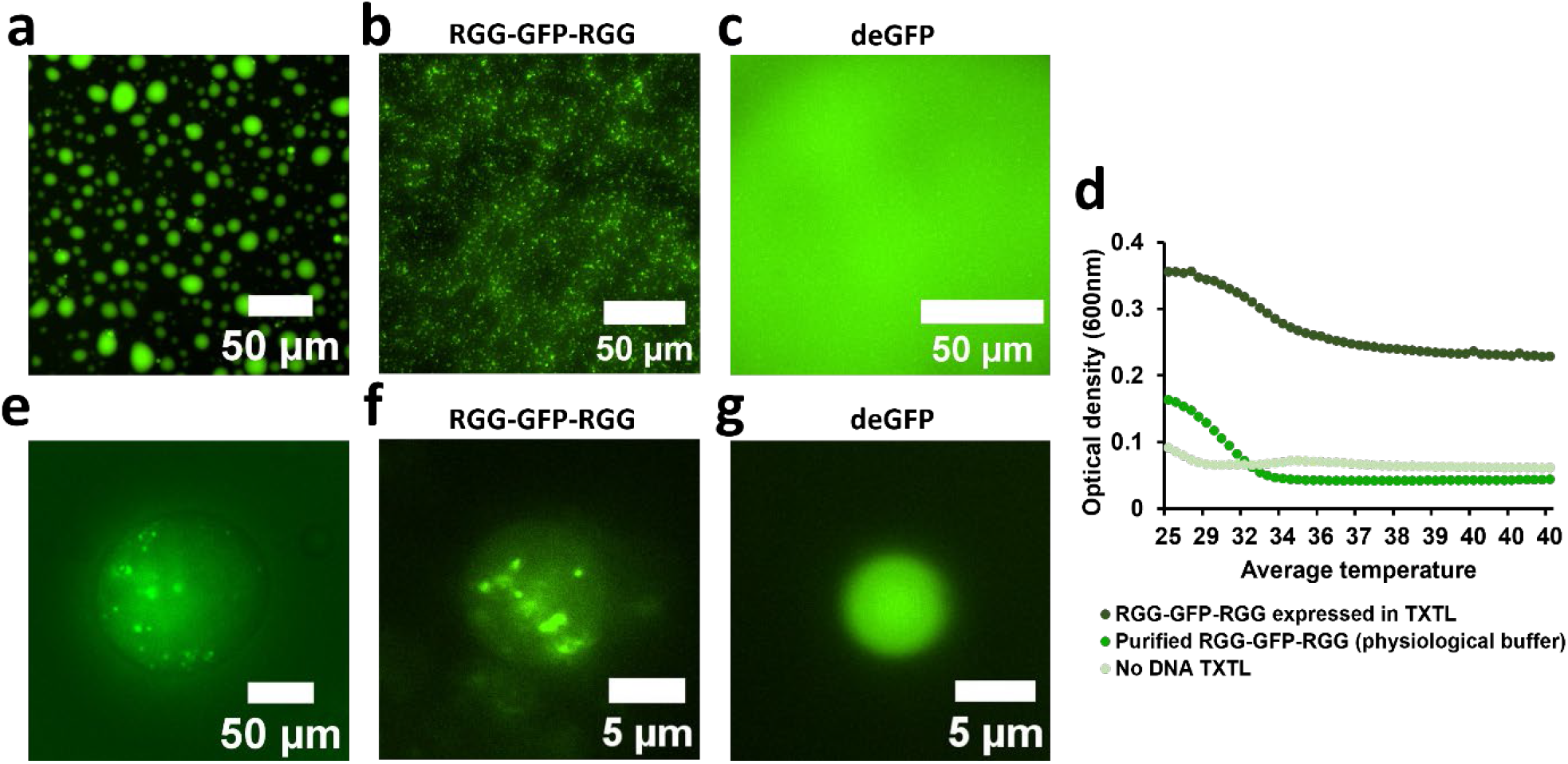
Phase separation of cell-free expressed RGG-GFP-RGG. **a**, RGG-GFP-RGG purified from *E. coli* (20 μM, in storage buffer). **b-c**, Phase separation of RGG-GFP-RGG expressed in TXTL reaction (**b**), compared to no phase separation observed in deGFP control (**c**). **d**, Turbidity assay depicts phase properties of RGG-GFP-RGG expressed by TXTL. **e**, RGG-GFP-RGG purified from *E. coli* forms condensates when encapsulated inside 1 mM POPC/1 mM cholesterol liposomes. **f-g**, RGG-GFP-RGG expressed from a plasmid in TXTL reaction forms condensates (**f**) when encapsulated inside 1 mM POPC/1 mM cholesterol liposomes; no condensates are observed in deGFP control (**g**).

The pET_RGG-GFP-RGG-His plasmid was expressed by TXTL, and a microplate fluorometer was used to assay fluorescence intensity of the GFP-tagged RGG protein over the course of the TXTL reaction to confirm expression (Supplementary Fig. 1). deGFP, a truncated version of eGFP which has been optimized for cell free expression, was also expressed as a positive control^33^. From this kinetic assay, we observed increased fluorescence of RGG-GFP-RGG, similar to the deGFP positive control. We also confirmed successful expression of RGG-GFP-RGG by western blot (Supplementary Fig. 2). The concentration of RGG-GFP-RGG expressed in TXTL was estimated to be approximately 5 μM (Supplementary Fig. 2). Together, these two assays indicate the protein can be expressed by TXTL. Since coacervate formation is dependent on salt, pH, and protein concentration, we next confirmed that when expressed by TXTL, RGG-GFP-RGG still forms coacervates. We imaged the reactions on a fluorescence microscope and observed protein-rich droplets in TXTL reactions expressing RGG-GFP-RGG, but not deGFP (Fig. 2b-c).

To characterize the temperature-dependent phase behavior of the TXTL-expressed protein, turbidity was measured at different temperatures using a microplate spectrophotometer (Fig. 2d, Supplementary Fig. 3). A TXTL reaction expressing RGG-GFP-RGG was prepared and incubated for 8 hours, and the resultant reaction (approximately 5 μM RGG-GFP-RGG concentration) was assayed for temperature-dependent turbidity, along with a TXTL reaction with no DNA template and RGG-GFP-RGG purified from E. coli (13.5 μM RGG-GFP-RGG concentration) in physiological buffer (150 mM NaCl, 6 mM Tris). Absorbance readings at OD600 were taken as the plate reader warmed from ambient temperature (∼25°C) to 40°C. It was previously reported that RGG-RGG, a protein containing two identical RGG domains to RGG-GFP-RGG but no GFP domain, in physiological buffer at 12 μM RGG domain concentration has a transition temperature around 40°C. In our assay, RGG-GFP-RGG had transition temperature around 35°C^27^. This lower transition temperature could be due to the presence of GFP, which is soluble. RGG-GFP-RGG expressed by TXTL had similar phase properties as the purified protein, with a critical temperature around 40°C, albeit the TXTL reaction was considerably more turbid compared to the no DNA control and purified RGG-GFP-RGG protein. This could be due to interaction of RGG-GFP-RGG with components present in the TXTL reaction mix. The phase behavior of RGG-GFP-RGG expressed by TXTL further suggests the protein is expressed and undergoes coacervation when expressed cell-free.

We wished to create a synthetic cell that is capable of producing its own microcompartments. To confirm the RGG-GFP-RGG protein forms coacervates when bound within a lipid membrane, we encapsulated the purified protein in liposomes. RGG-GFP-RGG was encapsulated in POPC/cholesterol liposomes using a water-in-oil emulsion method^34^. Phase separation of RGG-GFP-RGG was observed within the liposomes (Fig. 2e, Supplementary Fig. 4).

To engineer the synthetic cells expressing RGG-GFP-RGG condensates, TXTL reactions expressing the RGG-GFP-RGG plasmid were prepared and encapsulated in POPC/cholesterol liposomes. Following encapsulation, the liposomes were incubated at 30°C for expression and subsequently imaged using fluorescence microscopy. Fluorescent condensates were observed in the liposomes containing the RGG-GFP-RGG TXTL reaction (Fig. 2f), while none were observed in liposomes expressing deGFP (Fig. 2g), showing that cell-free expression of RGG-GFP-RGG within liposomes results in the formation of condensates. Sufficient protein concentration of RGG-GFP-RGG was necessary for condensate formation within liposomes. The morphology of the condensates appears distinct from that of multilamellar and multivesicular vesicles (Supplementary Fig. 5)

### Creating RNA-binding membraneless organelles in synthetic cells

After verifying we can create condensates within synthetic cells, we then sought to functionalize these microcompartments. Membraneless organelles are known to play a role in controlling gene expression through mRNA regulation, and RGG-domains are known RNA binding domains, so we wished to exploit RNA-binding of the cell-free expressed RGG-GFP-RGG coacervates in synthetic cells^21,23–25^.

RNA binding to RGG domains is well known and has been characterized previously by electrophoretic motility shift assays (EMSA)^35–37^. To confirm binding of labeled RNA in our system, we used EMSA to observe interactions between RGG-GFP-RGG and RNA. A short (36 nucleotide) RNA encoding for a fragment of deGFP was transcribed, incubated with RGG-GFP-RGG, and ran on an agarose gel (Fig. 3a). The intensity of the band of deGFP RNA that migrated on the gel was quantified (Fig. 3b). The unbound RNA in the sample with RGG-GFP-RGG present had a 27% decrease in intensity compared to the RNA-only control, indicating that RNA remained bound to RGG-GFP-RGG and did not migrate down the gel. Smearing was also noted in lanes with RGG-GFP-RGG, suggesting that bound nucleic acid carried over from purification.

**Fig. 3:**
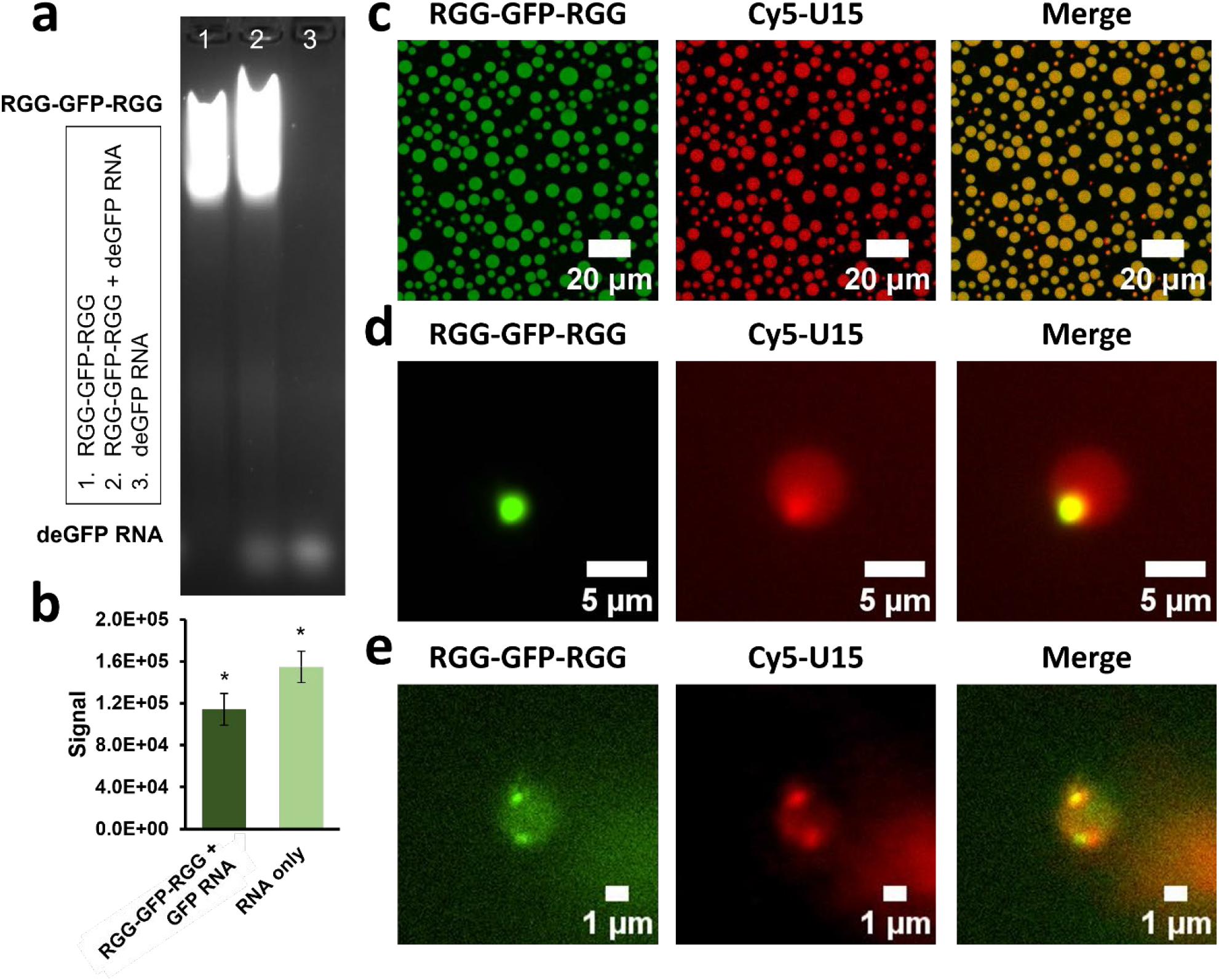
RNA sequestration by RGG-GFP-RGG membraneless organelles. **a**, RGG-GFP-RGG (13.5 μM) and deGFP RNA (∼20 μg) on 1% agarose gel stained with SYBR-Safe. **b**, Gel quantification of migrated deGFP RNA bands in (**a**). Binding of RNA to RGG-GFP-RGG reduced intensity of migrated RNA band. Error bars signify SEM (n = 3) as determined by t-test. *\*p < 0.05*. **c**, Confocal view of RGG-GFP-RGG coacervates binding Cy5-U15 RNA. **d**, RGG-GFP-RGG and Cy5-U15 encapsulated in POPC/cholesterol liposomes binds Cy5-U15. **e**, A TXTL reaction expressing RGG-GPF-RGG was encapsulated with Cy5-U15 in liposomes. Cy5-U15 localization to the RGG-GFP-RGG droplets is observed.

Further, Cy5-labelled polyU(15) RNA (Cy5-U15) was used to visualize RNA sequestration by RGG-GFP-RGG. Cy5-U15 was added to purified RGG-GFP-RGG and imaged, and clear localization of Cy5-U15 to RGG-GFP-RGG was observed (Fig 3c). Cy5-U15 and RGG-GFP-RGG were then encapsulated in POPC/cholesterol liposomes, where localization of Cy5-U15 RNA to the condensates was seen once again (Fig. 3d). We then wished to confirm sequestration of Cy5-U15 RNA by coacervates expressed in cell-free reactions. TXTL reactions expressing RGG-GFP-RGG were prepared, and the reaction and Cy5-U15 were encapsulated in liposomes and incubated. After expression, RGG-GFP-RGG condensates were visible in the liposomes, with Cy5-U15 localized to the condensates (Fig. 3e). Cy5-U15 localization to deGFP expressed by TXTL was not observed (Supplementary Fig. 6, Supplementary Fig. 7).

### RNA sequestration by RGG-GFP-RGG condensates affects downstream protein expression

Since coacervates can sequester RNA or other molecules necessary for TXTL expression and create a molecularly crowded environment, we first verified that other proteins can be expressed in the presence of RGG-GFP-RGG coacervates^38^. TXTL reactions expressing deGFP were prepared, with the addition of increasing concentrations of purified RGG-GFP-RGG protein. The resultant expression of deGFP was then characterized by western blot (Fig. 4a). The western blot showed that deGFP was expressed by TXTL in the presence of the IDP. Interestingly, there is an inverse relationship between the amount of RGG-GFP-RGG present and the level of deGFP expression: as more IDP is added to the reaction, deGFP expression decreases. Our hypothesis is that RGG-GFP-RGG binds transcriptional RNA during cell-free expression, resulting in lessened protein expression.

**Fig. 4:**
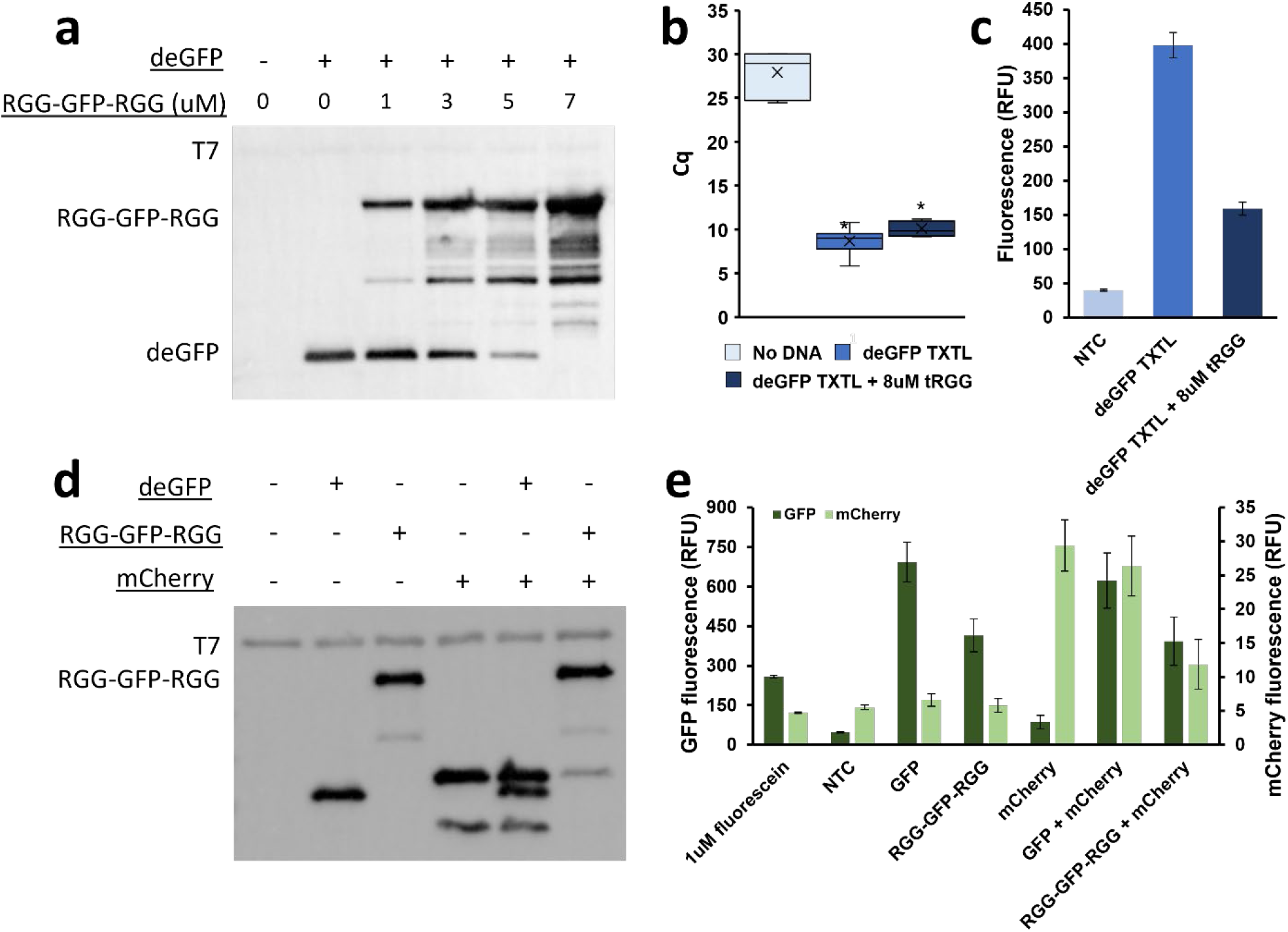
Sequestration of transcriptional mRNA by RGG proteins reduces protein expression in TXTL. **a**, TXTL reactions expressing deGFP were prepared and supplemented with increasing concentrations of RGG-GFP-RGG. RGG-GFP-RGG reduces deGFP expression in concentration dependent manner **b**, RT-qPCR was performed targeting deGFP in TXTL reactions with tRGG added. Prior to RT, reactions were spun down to remove tRGG and bound mRNA and the supernatant was taken for RT and subsequent qPCR. Reactions with tRGG present had elevated Cq values, indicating mRNA abundance was reduced when tRGG was removed. Error bars signify SEM (separate TXTL reactions were considered biological replicates (n=3); each biological replicate underwent qPCR in triplicate (n=3)) as determined by t-test *\*p < 0.05*. **c**, deGFP fluorescence in TXTL reactions with tRGG added was reduced. **d-e**, TXTL reactions expressing multiple plasmids were prepared and analyzed by western blot (**d**) and spectrofluorometer (**e**). Two-plasmid TXTL reactions expressing GFP and mCherry had fluorescence values slightly reduced compared to controls; TXTL reactions expressing both RGG-GFP-RGG and mCherry showed expected fluorescence values for RGG-GFP-RGG, while mCherry fluorescence was significantly reduced.

To confirm that RGG-GFP-RGG reduces protein expression by sequestering RNA, we used quantitative reverse transcription PCR (RT-qPCR) to quantify mRNA abundance in TXTL reactions when RGG protein is present (Fig 4b). TXTL reactions expressing deGFP were prepared and non-fluorescent RGG-RGG was added. RGG-RGG contains the same RGG domains as RGG-GFP-RGG, but contains no GFP and was used instead of RGG-GFP-RGG to reduce potential artifacts from using the GFP-tagged protein during qPCR. Prior to RT, the samples were spun down to pellet the condensates and the supernatant containing transcriptional mRNA was used for further reactions. Flourescence readings of GFP from the TXTL reactions were also taken (Fig. 4c). Both mRNA abundance and fluorescence were depleted in TXTL reactions with RGG-RGG added, suggesting that RGG binds mRNA from transcription, thereby reducing the availability of mRNA to be translated into deGFP.

To see if this effect was seen when expressing RGG-GFP-RGG cell-free, we set up TXTLs with two plasmid templates, expressing both RGG-GFP-RGG and mCherry, another fluorescent protein, in one reaction (Fig. 4). Our hypothesis was that the expressed RGG-GFP-RGG protein would phase separate to form model membraneless organelles that would in turn sequester transcriptional mRNA needed for mCherry expression, reducing mCherry fluorescence signal. This was indeed observed, with mCherry fluorescence signal decreased by 55% when expressed with RGG-GFP-RGG as compared to co-expression with deGFP (Fig. 4e). Of note, mCherry has a lower overall brightness compared to GFP resulting in lower average RFU values^39^. To confirm that the decrease in fluorescence was not due to fluorescence artifacts from the presence of the RGG-GFP-RGG droplets, protein expression was also measured by western blot (Fig. 4d). The two-plasmid TXTL of RGG-GFP-RGG and mCherry showed decreased mCherry band intensity on western blot compared to the positive control TXTL expressing both GFP and mCherry, indicating that mCherry expression was significantly hindered by the presence of RGG-GFP-RGG droplets. Together, this demonstrates that when co-expressed using TXTL, RGG-GFP-RGG membraneless organelles can modulate cell-free expression of another plasmid.

## Discussion

Membraneless organelles have garnered increased interest in recent years as we learn more about their functions in natural cells, including their role in regulation of protein expression. Cells respond to stress by remodeling their transcriptome through transcription and degradation of mRNA. Membraneless organelles like P-bodies and stress granules store mRNA, affecting mRNA localization, translation, and degradation^40,41^. We aimed to design a system where the compartments inside synthetic cells can bind RNAs, paving the way for a platform for synthetic cells to regulate their own gene expression as these membraneless organelles do in living cells.

Increasing the complexity of and creating versatile tools for synthetic cells is a major pursuit in synthetic biology. In this work, we demonstrated synthetic cells expressing the coacervate-forming protein RGG-GFP-RGG to function as artificial membraneless organelles. Those organelles were shown to sequester mRNA from TXTL reactions and reduce protein expression of plasmid-encoded proteins. Cellular organelles allow for certain biological processes to occur in a specialized environment, separated from the outside influences of the cell. These specialized compartments are critical for the numerous and complex functions of natural cells. Building organelles within synthetic cells not only increases the capabilities of synthetic cells, but also provides insight on spatiotemporal organization of natural biological cells and their origins.

## Methods

### Plasmid and oligonucleotide constructs

Plasmid information can be found in Supplementary Table 1. Oligonucleotide sequences can be found in Supplementary Table 2. pET_RGG-GFP-RGG and pET_RGG-RGG were gifts from Matthew Good, Daniel Hammer, and Benjamin Schuster (Addgene plasmids #124948, #124941). For transcription of RNA transcripts, corresponding DNA oligonucleotides were ordered from Integrated DNA Technologies (IDT). Cyanine5-labeled U(15) RNA (Cy5-U15) was custom purchased from Sigma-Aldrich.

### Purification of RGG-GFP-RGG and RGG-RGG from E. coli

RGG-GFP-RGG and RGG-RGG were purified according to the methods previously described^5^. Plasmids were transformed in BL21(DE3) *E. coli* for protein expression. Cultures were grown in Terrific Broth (TB) media at 37°C while shaking at 250RPM until reaching OD600 = 0.6. Cultures were then induced with 1 mM isopropyl β-d-1-thiogalactopyranoside (IPTG) and incubated at 18°C, 225 RPM overnight for protein expression. The bacterial pellets were resuspended in high salt lysis buffer (500 mM NaCl, 20 mM Tris, 20 mM imidazole, 1mM PMSF, pH 7.5) and sonicated. To prevent phase separation, lysates were centrifuged at 15,000g for 30 minutes at 40C. Proteins were washed (500 mM NaCl, 20 mM Tris, 20 mM imidazole, pH 7.5), purified using Nickel NTA agarose beads (Catalog No. H-350-5, GoldBio), and eluted (500 mM NaCl, 20 mM Tris, 500 mM imidazole, pH 7.5). Finally, proteins were dialyzed overnight using a 7 or 10 kDa molecular weight cut-off Slide-A-Lyzer dialysis cassette (Catalog No. 66380, Thermo Scientific) in high salt buffer (500 mM NaCl and 20 mM Tris, pH 7.5) at 42°C to prevent phase separation. The dialyzed protein was aliquoted, flash frozen in liquid nitrogen, and stored at -80C.

### Cell-free protein expression (TXTL) and quantification

Cell-free protein expression of RGG-GFP-RGG was carried out using a T7-based TXTL system adopted from the Noireaux lab and using methods described previously^34^. Cell extract was prepared from Rosetta2 *E. coli* by methods described previously^42^. TXTL reactions contain 10 nM DNA template, 12 mM magnesium glutamate, 140 mM potassium glutamate, 1 mM DTT, 1.5 μM T7 RNA polymerase, 0.4 U/μL Murine RNase Inhibitor, 1× energy mix, 1× amino acid mix, and cell free extract. The 10× energy mix is composed of the following: 500 mM HEPES, pH 8, 15 mM ATP, 15 mM GTP, 9 mM CTP, 9 mM UTP, 2 mg/mL E. coli tRNA, 0.68 mM Folinic Acid, 3.3 mM NAD, 2.6 mM Coenzyme-A, 15 mM Spermidine, 40 mM Sodium Oxalate, 7.5 mM cAMP, and 300 mM 3-PGA. The 10× amino acid mix is composed of 17 mM of the following amino acids: alanine, arginine, asparagine, aspartic acid, cysteine, glutamic acid, glutamate, glycine, histidine, isoleucine, leucine, lysine, methionine, phenylalanine, proline, and serine.

For analysis of TXTL reactions using fluorescence, signal quantification was measured in 384-well plates (Catalog No. 09-761-85, Corning) using a microplate fluorometer (SpectraMax GeminiXS). Relative protein expression was estimated using western blot (Bio-Rad ChemiDoc).

### Cell-free transcription

Short linear templates were used for expression of RNA transcripts. Sense and anti-sense oligos containing a T7Max promoter sequence and a short fragment (36 nucleotides) of the coding sequence for deGFP were synthesized (Integrated DNA Technologies) and annealed in a thermocycler. A master mix of transcription reagents was prepared on ice containing DNA template, 1X homemade NEB Buffer, 8 mM GTP, 4 mM A/C/UTP, 0.005X phosphatase 25 ng/μ L, 1 μM T7 RNAP, and 0.4 U/μL Murine RNAse Inhibitor (Catalog No. M0314S, New England BioLabs). Reactions were incubated for 8 hours at 37°C in a thermocycler. RNA was purified using a Monarch RNA Cleanup Kit (500 μg) (Catalog No. T2050L, New England BioLabs) and concentration was measured using a NanoDrop ND-1000 UV-Vis Spectrophotometer (Thermo Scientific).

### Turbidity assay

Phase properties of the protein were determined by measuring optical density (OD) at 600 nm using a microplate spectrophotometer (SpectraMax 340PC384). 100 μL TXTL reactions expressing RGG-GFP-RGG were prepared as well as a no DNA control TXTL reactions. Purified RGG-GFP-RGG in physiological buffer was also assayed. Samples were prepared in a clear 96-well microplate and readings were taken every 30 seconds as the instrument heated from ambient temperature (∼25°C) to 40°C.

### Liposome preparation and encapsulation

Thin films were prepared by dissolving 1mM 1-palmitoyl-2-oleoyl-sn-glycero-3-PC (POPC) and 1 mM cholesterol (Catalog No. 15102, Catalog No. 9003100, Cayman Chemical) in chloroform or dichloromethane in glass vials. Vials were left uncapped overnight in a fume hood to allow for the solvent to evaporate and thin films to form. Thin films were resuspended in mineral oil (Catalog No. 1632129, Bio-Rad) to create the lipid-in-oil solution. Internal contents were encapsulated in liposomes using water-in-oil emulsion method described previously^34^. Nile Red was used to visualize liposome membranes (Catalog No. N1142, Thermo Scientific).

### Electrophoretic mobility shift assay (EMSA)

deGFP RNA binding to RGG-GFP-RGG was visualized by agarose gel electrophoresis. ∼15 μg of purified deGFP RNA was added to 13.5 μM RGG-GFP-RGG in 150 mM NaCl and 6 μM Tris and loaded into a 1% agarose gel stained with 1X SYBR Safe DNA Gel Stain (Catalog No. S33102, Thermo Fisher). The mRNA was separated from the protein at 125 V for 45 minutes in 1X TAE buffer and gels were imaged (Aplegen Omega Lum G). The unbound RNA bands were quantified using Fiji.

### Measuring mRNA abundance with quantitative reverse transcription PCR (RT-qPCR)

TXTL and transcription reactions expressing deGFP were prepared in triplicate with either 8 μM RGG-RGG added or the corresponding volume of storage buffer. After incubation, TXTL and transcription reactions were centrifuged at 3200 g for 10 minutes to pellet RGG-RGG and bound mRNA. The supernatant was removed and treated with DpnI to degrade template plasmid DNA (Catalog No. R0176S, New England BioLabs Inc.). The mixture was incubated at 37°C for 15 minutes (T100 Thermal Cycler, Bio-Rad). The enzyme and the expressed proteins were inactivated by adding 15mM EDTA (Catalog No. E9884, Sigma-Aldrich) at 75°C for 10 minutes. The denatured proteins were pelleted through centrifugation at 3200 g for 2 minutes.

For the reverse transcription reaction, 4.5 μl of digested sample was added to 0.5μl of reverse primer (Integrated DNA Technologies), 0.5 μl of 10mM dNTP mix (Catalog No. CB4420-2, Denville Scientific), and 1 μl water. The reaction was incubated at 65°C for 1 minute, then put on ice for at least 5 minutes. The digested sample was then added to 0.5 μl of Invitrogen SuperScript IV Reverse Transcriptase (Catalog No. 18090200, Thermo Fisher), 2 μl of 5x Superscript IV Buffer, 0.5 μl of 100mM DTT, and 0.5 μl of RNase Inhibitor, Murine (Catalog No. M0314S, New England BioLabs Inc.). The full reaction was then incubated at 52°C for 10 minutes, and the reverse transcriptase was inactivated at 80°C for 10 minutes.

The qPCR reaction was prepared in triplicate with 2 μl of the resultant cDNA from the reverse transcription, 2 μl of 10 μM forward and reverse qPCR primers (Integrated DNA Technologies), 11.25 μl OneTaq Hot Start 2X Master Mix with Standard Buffer (Catalog No. M0484, New England BioLabs Inc.),1.25 μl Chai Green Dye 20X (Catalog No. R01200, Chai Bio), and 6.5 μl of nuclease-free water. The RT-qPCR was completed using CFX96 Touch Real-Time PCR Detection System (Bio-Rad) with the following thermocycling program: 1 cycle of 30 second denaturation at 94°C, 30 cycles of 15 second denaturation at 94°C, 15 second annealing at 54°C, 1 minute extension at 68°C, and 1 cycle of 5 minutes final extension at 68°C. The amplification curves and Cq values were calculated through the CFX Maestro software. The averages across 3 biological and technical replicates were calculated, replacing non-detected values with 30 (the maximum cycle number).

### Fluorescence microscopy

Fluorescence and brightfield images were taken using a Nikon Eclipse Ti microscope and confocal images were taken using a Leica TCS SP5 microscope. Image brightness, contrast, and channel color were adjusted and scale bars were added using Fiji (ImageJ).

## Acknowledgments

All authors were supported by the NSF award 1844313 RoL: RAISE: DESYN-C3: Engineering multi-compartmentalised synthetic minimal cells. AR, AC and KPA were supported by the Alfred P. Sloan Foundation grant G-2022-19420, NSF award 2227578 Moving Millions of Droplets at Megahertz Speeds. AR was supported by NIH Biotechnology Training Grant 5T32GM008347-29 and 5T32GM008347-30. Figures created with BioRender.com.

## Supplementary Information (SI)

**Supplementary Table 1.** Plasmid information

| Name | Source |
| --- | --- |
| pET_RGG-GFP-RGG (His) | Schuster BS, Reed EH, Parthasarathy R, Jahnke CN, Caldwell RM, Bermudez JG, Ramage H, Good MC, Hammer DA. Nat Commun. 2018 Jul 30;9(1):2985. doi: 10.1038/s41467-018-05403-1. |
| pET_RGG-RGG (His) | Schuster BS, Reed EH, Parthasarathy R, Jahnke CN, Caldwell RM, Bermudez JG, Ramage H, Good MC, Hammer DA. Nat Commun. 2018 Jul 30;9(1):2985. doi: 10.1038/s41467-018-05403-1. |
| T7Max-UTR1-deGFP-8xHis-T500 | Deich, C., Cash, B., Sato, W. *et al.* T7Max transcription system. *J Biol Eng* **17**, 4 (2023). doi: 10.1186/s13036-023-00323-1 |
| T7Max-UTR1-mCherry-8xHis-T500 | Deich, C., Cash, B., Sato, W. *et al.* T7Max transcription system. *J Biol Eng* **17**, 4 (2023). doi: 10.1186/s13036-023-00323-1 |

**Supplementary Table 2.**
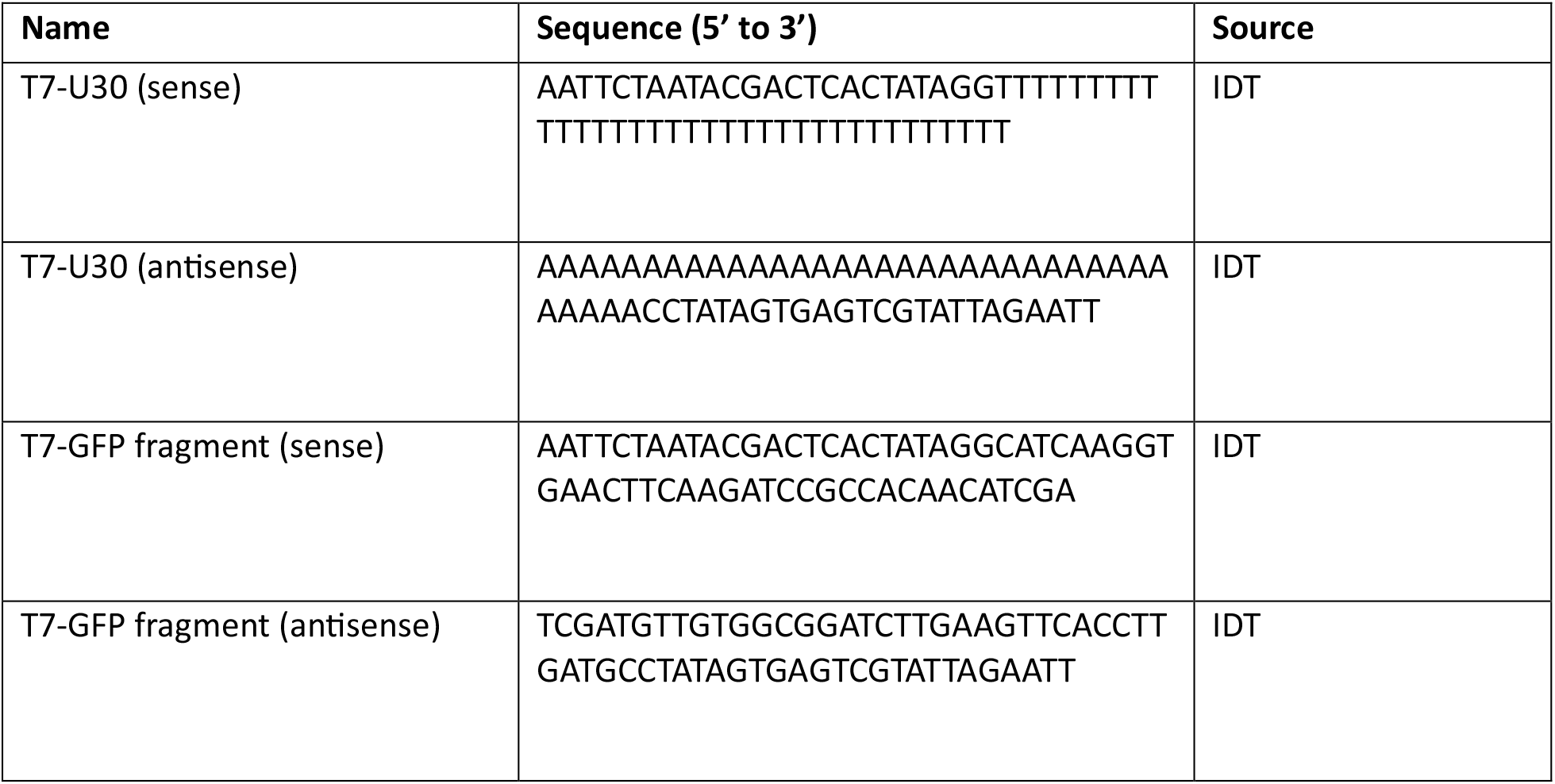
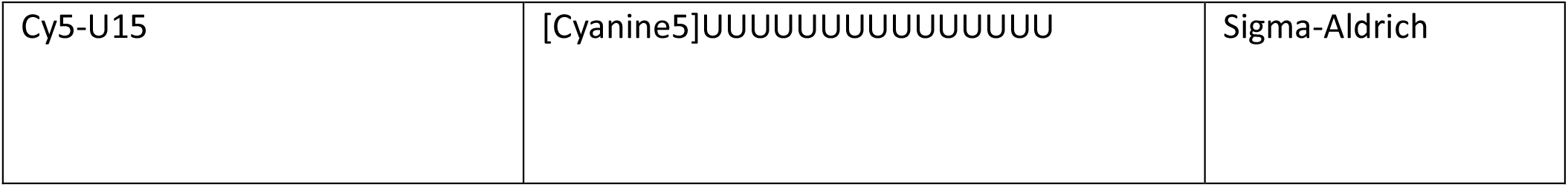
Oligonucleotide sequences

**Supplementary Fig. 1:**
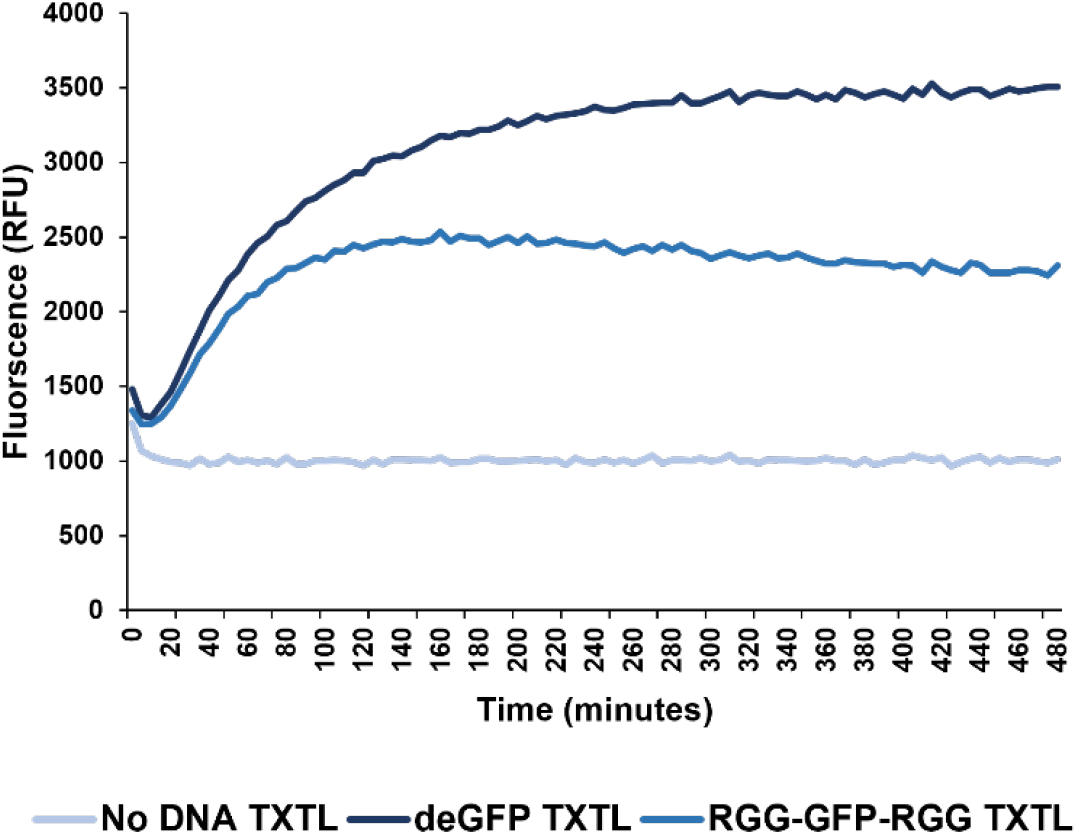
Kinetic fluorescence data of RGG-GFP-RGG TXTL reaction. TXTL reactions expressing RGG-GFP-RGG, a positive control (deGFP), and a no DNA control were prepared in 364-well microplates and fluorescence readings were taken for 8 hours (480 minutes) on a microplate fluorometer (SpectraMax GeminiXS). Relative fluorescence of RGG-GFP-RGG was significant compared to the no DNA TXTL reaction, suggesting protein expression.

**Supplementary Fig. 2:**
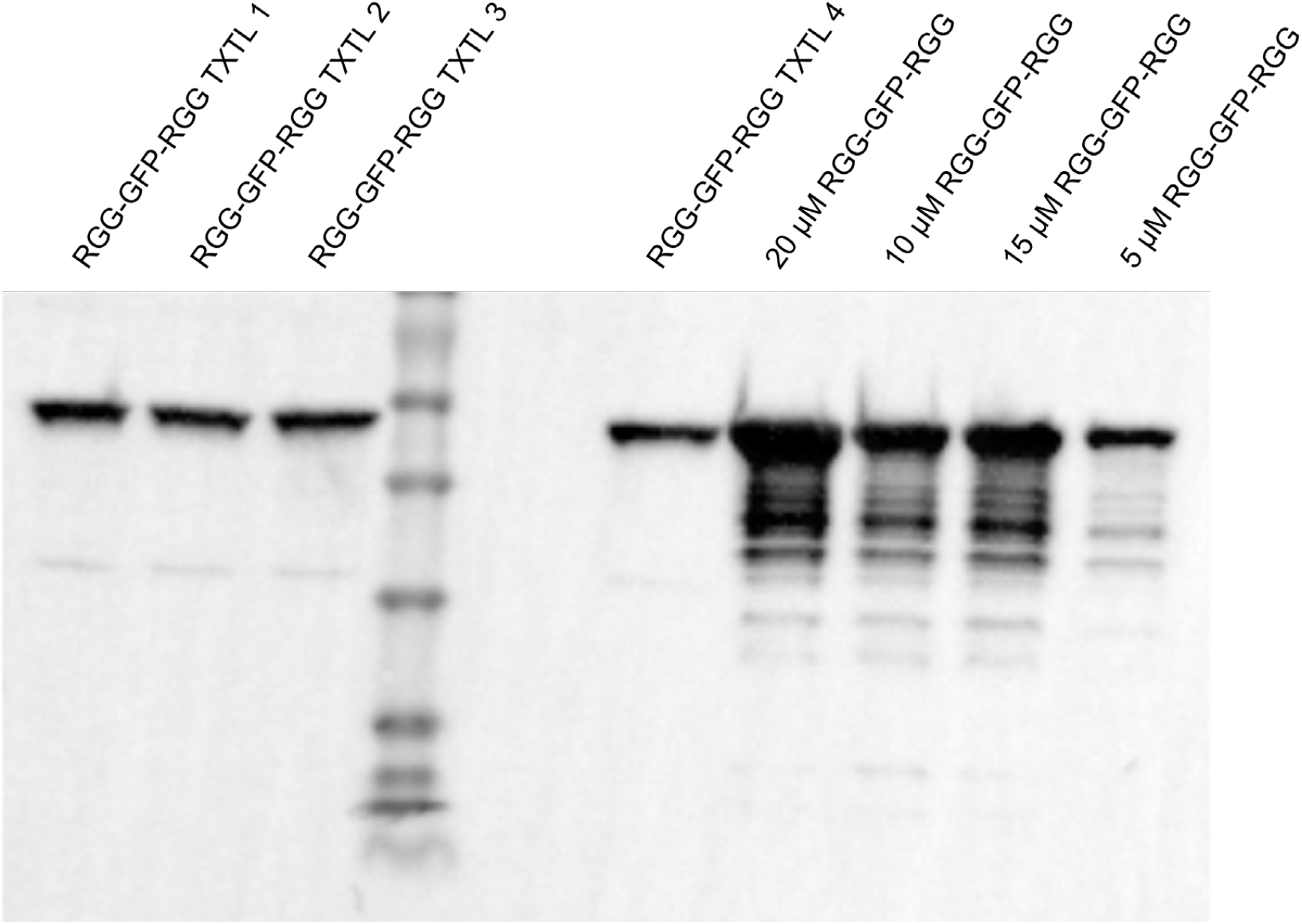
Western blot for estimation of RGG-GFP-RGG protein concentration from TXTL expression. Four TXTL reactions expressing RGG-GFP-RGG were prepared and blotted alongside known concentrations of RGG-GFP-RGG produced by overexpression in *E. coli* and purification as described in Methods. RGG-GFP-RGG concentration is estimated to be around 5 μM.

**Supplementary Fig. 3:**
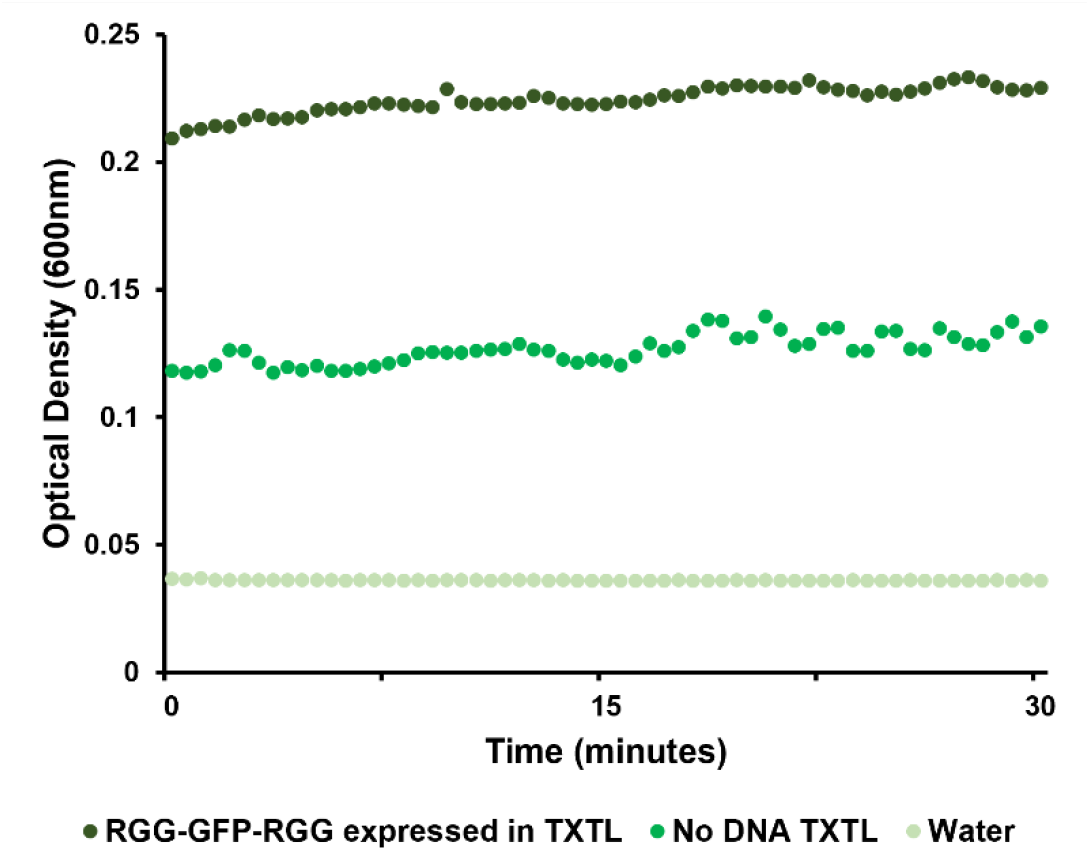
Turbidity assay for TXTL-expressed RGG-GFP-RGG at 26°C. RGG-GFP-RGG was expressed by TXTL and turbidity was measured close to room temperature, confirming the protein solutions remain turbid over time prior to temperature ramp experiment (Fig. 2d).

**Supplementary Fig. 4:**
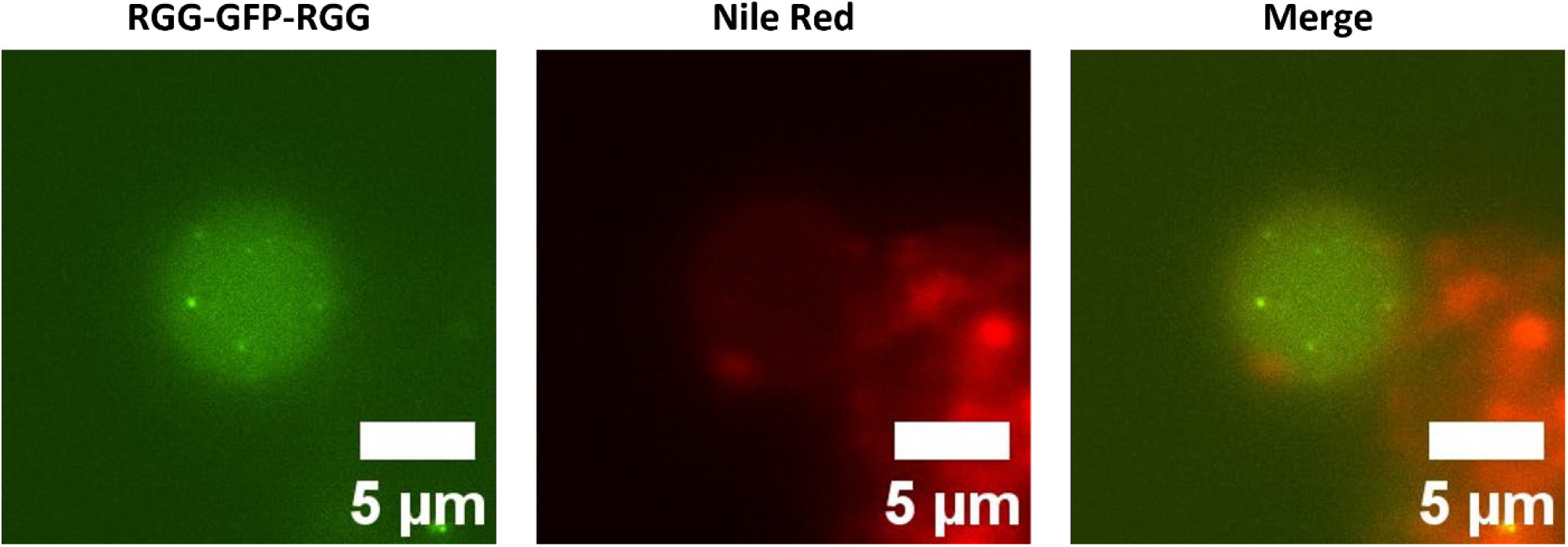
Phase separation of cell-free expressed RGG-GFP-RGG inside Nile Red stained liposomes. Phase separation of RGG-GFP-RGG is observed inside of liposomes; the liposome membrane is labeled with Nile Red.

**Supplementary Fig. 5:**
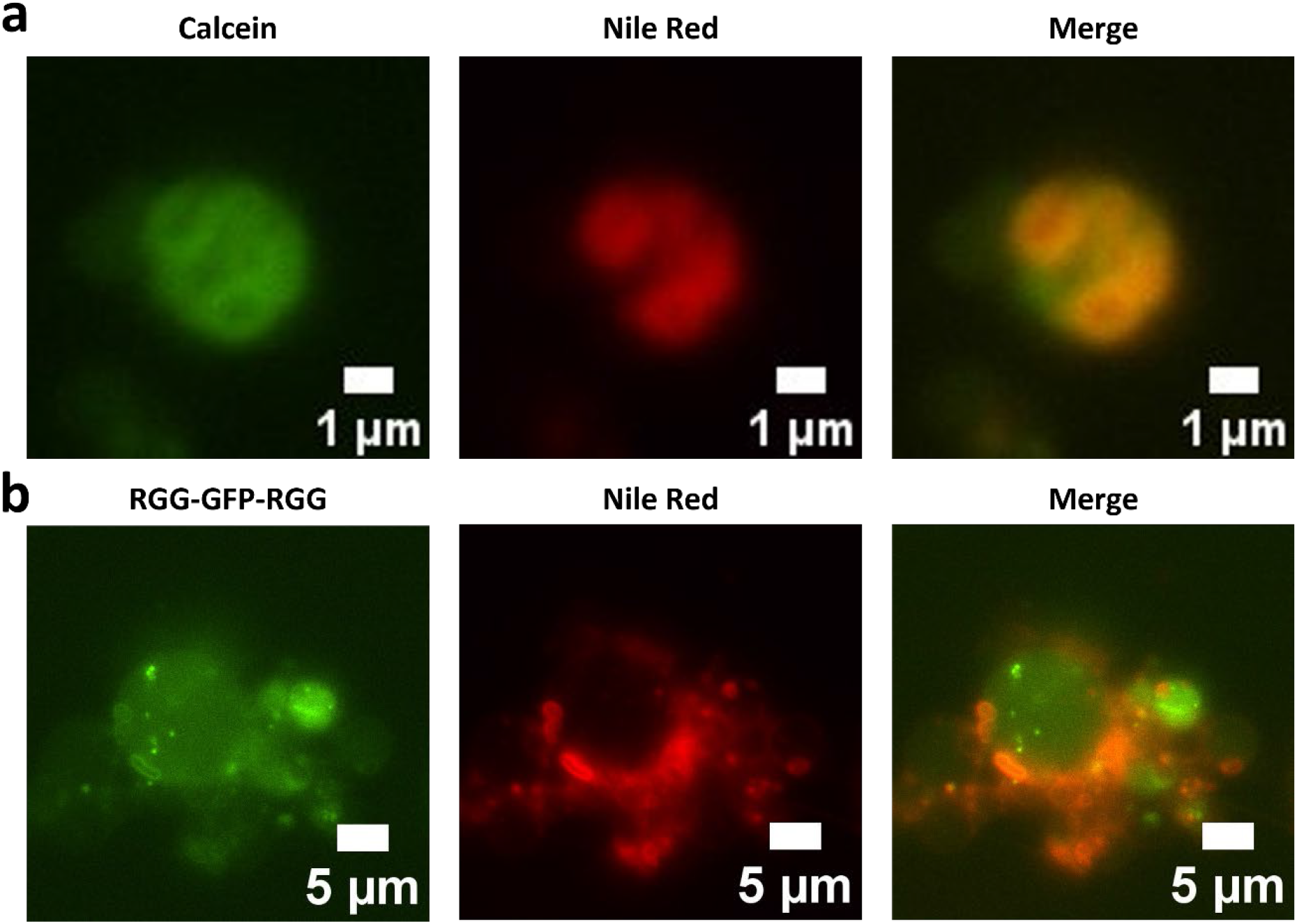
Microscopy imaging of multivesicular liposomes. **a**, Calcein dye was encapsulated in POPC/cholesterol liposomes; liposome membrane is stained with Nile Red. Multiple vesicles are encapsulated inside of a larger vesicle. **b**, TXTL reactions expressing RGG-GFP-RGG were encapsulated inside of liposomes and membranes stained with Nile Red. RGG-GFP-RGG coacervates are visible, as well as additional vesicles encapsulated inside of a larger liposome.

**Supplementary Fig. 6:**
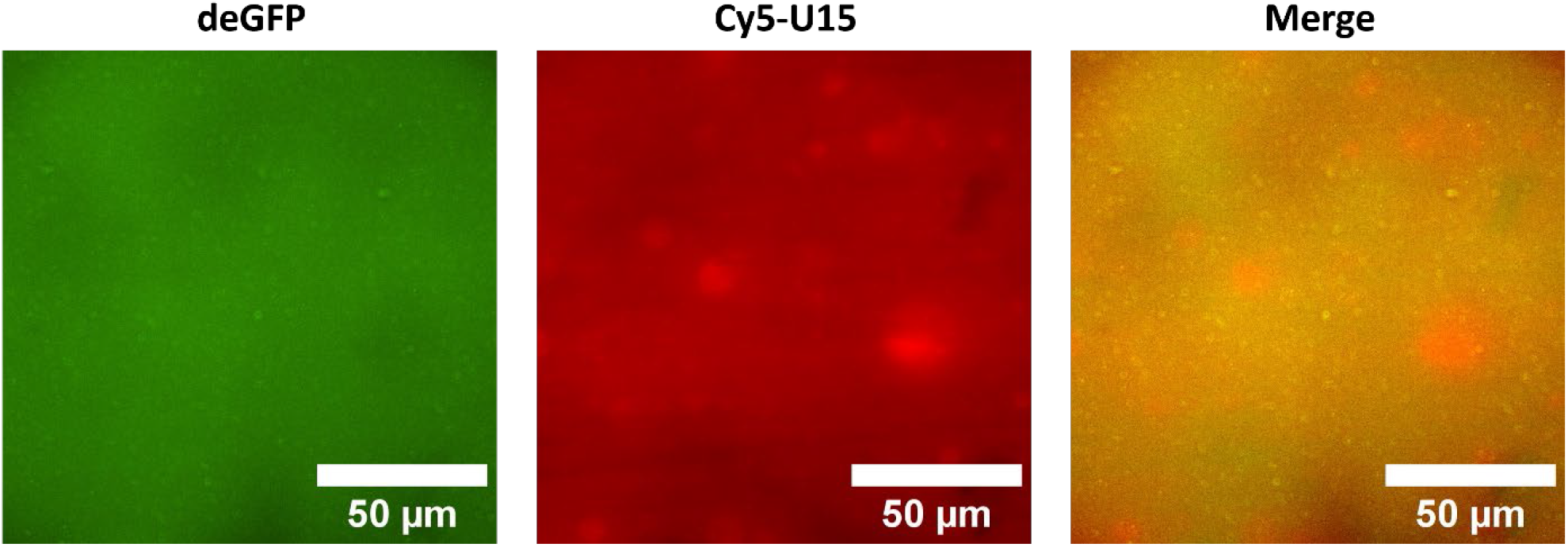
Microscopy imaging of deGFP expressed by TXTL with Cy5-U15 RNA. TXTL reactions expressing deGFP were prepared with Cy5-U15 RNA and were imaged following incubation. No clear binding of Cy5-U15 RNA to deGFP was observed.

**Supplementary Fig. 7:**
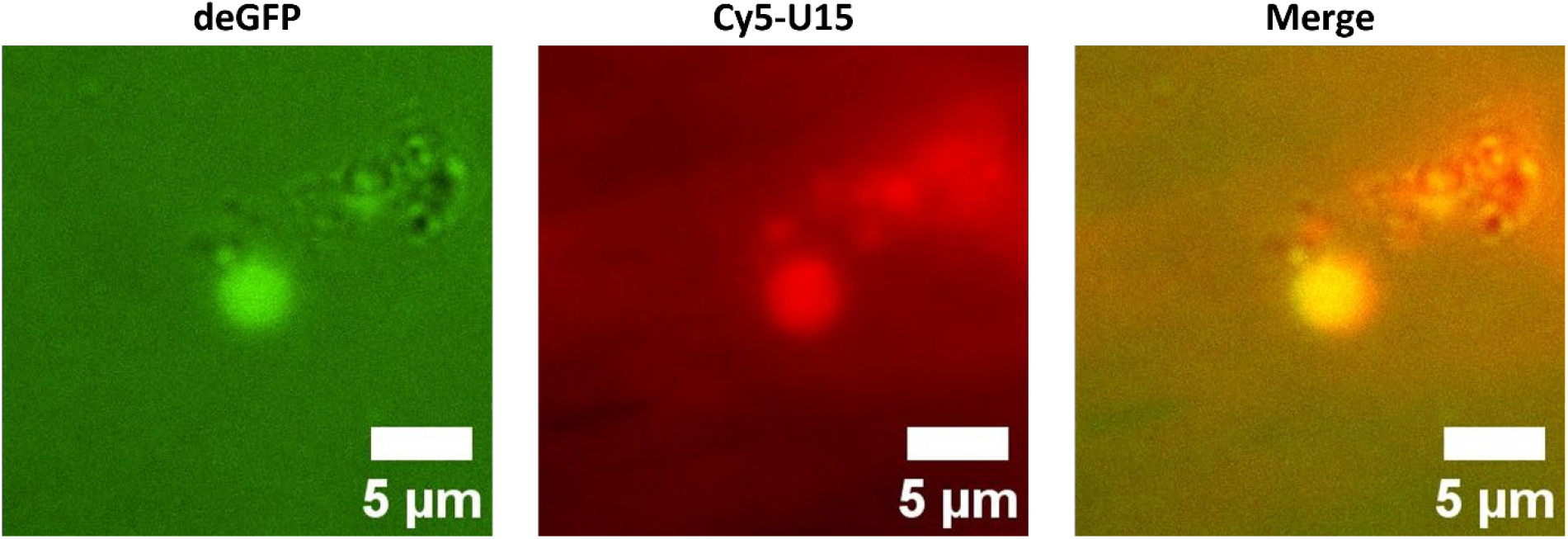
Microscopy imaging of deGFP TXTL with Cy5-U15 RNA inside of liposomes. No coacervate structures were observed, nor binding of Cy5-U15 RNA to deGFP.

**Supplementary Video 1**

RGG-GFP-RGG coacervates expressed by TXTL encapsulated in a POPC/cholesterol liposome.

